# A novel and simple heat-based method eliminates the highly detrimental effect of xylene deparaffinization on acid-fast stains

**DOI:** 10.1101/2022.11.29.518155

**Authors:** Pedro F. Marinho, Soraia L. Vieira, Tânia G. Carvalho, Maria C. Peleteiro, Thomas Hanscheid

**Affiliations:** Instituto de Microbiologia, Faculdade de Medicina, Universidade de Lisboa, 1649-028 Lisboa, Portugal; Instituto de Medicina Molecular João Lobo Antunes, Edifício Egas Moniz, Avenida Professor Egas Moniz, 1649-028 Lisboa, Portugal; Fundação Champalimaud, Avenida Brasília, 1400-038 Lisboa, Portugal; Centro de Investigação Interdisciplinar em Sanidade Animal, Faculdade de Medicina Veterinária, Universidade de Lisboa, Av. da Universidade Técnica, 1300-477 Lisboa, Portugal

**Keywords:** acid-fast-stain, xylene-deparaffinization, histology, mycobacteria, novel method

## Abstract

**Objectives:** Histopathology is an important method for the diagnosis of extrapulmonary tuberculosis, yet tissue sections are often negative for mycobacteria after acid-fast-staining (AFS). This study investigated the mechanism of AFS and the detrimental effect of histological processing, in particular xylene deparaffinization, on AFS staining and mycobacterial detection.

**Methods:** The target of the fluorescent Auramine O (AuO) AFS was investigated using triple staining with DNA and RNA specific dyes. The effect of xylene deparaffinization on acid-fastness of mycobacteria in cultures or tissue sections was studied using AuO fluorescence as a quantitative marker. The xylene method was compared to a novel, solvent-free projected-hot-air-deparaffinization (PHAD).

**Results:** Colocalization of AuO with DNA/RNA stains suggests that intracellular nucleic acids are the true target of AFS, producing highly specific patterns. Xylene reduces mycobacterial fluorescence significantly (p<0,0001, moderate effect size: *r* = 0.33). PHAD yielded significantly higher fluorescence than xylene deparaffinization in tissues (p<0.0001, large effect size: *r* = 0.85).

**Conclusions:** AuO can be applied for nucleic-acid-staining of mycobacteria in tissues producing typical beaded patterns. AFS depends crucially on the integrity of the mycobacterial cell-wall, which seems to be damaged by xylene. A solvent-free tissue deparaffinization method has the potential to increase mycobacterial detection significantly.

**Key Points:** Acid-fast-stains for mycobacterial detection have a notoriously poor yield, due to misconceptions of the underlying staining principle, but mainly because of the possible detrimental effect of xylene.

The yield of acid-fast-stains for mycobacterial detection in histology, including fluorescent stains, can be significantly increased by eliminating solvents (xylene) in the deparaffinization step.

Acid-fast-stains depend on bacterial integrity because they target nucleic acids. A simple hot air deparaffinization is significantly superior to xylene, avoiding damage to the lipid-rich cell wall.

## Introduction

Tuberculosis (TB) remains a serious infectious disease, with some 10 million cases and, despite effective treatment, 1.5 million deaths per year.^1^ Pulmonary TB is usually diagnosed reliably by microbiological methods, employing light or fluorescent microscopy and acid-fast stains (AFS), culture, or nucleic acid amplification tests (NAAT).^2^ Contrary to this, extrapulmonary tuberculosis (EPTB), which afflicted one in six patients (16%) in 2019, is a paucibacillary condition which often requires biopsies.^1,3^ Bacteriological diagnosis, is difficult as illustrated by the 2021 UK TB report, where the majority of EPTB cases (56%) were culture-negative^3,4^ and only 4% of these had a positive NAAT. However, 14% of these were positive on histological examination, ^4^ demonstrating the importance of histology for the etiological diagnosis of tuberculosis. However, while histology often reveals TB-like tissue lesions, such as granulomas, AFS stained sections, needed for a definitive diagnosis, are often negative for mycobacteria.^3,5,6^ Using AFS on deparaffinized histology sections is well-known for its poor yield which is attributed to the low number of mycobacteria present in the tissue.^3,5,7–9^ However, could it be that histologic processing and the staining procedures themselves cause the unreliable detection of mycobacteria?

Mycolic acids (MA) of mycobacteria are often reported to be the target of AFS, for both carbol-fuchsin used in light-microscopy and Auramine O (AuO) used in fluorescent microscopy.^10–12^ Yet, both are recognized nucleic acid stains, ^13–15^ which seems to imply that the AFS depends on the integrity of the MA layer to prevent the removal of the primary dye during acid-decolorization.

Histological processing of tissue is usually based on paraffin embedded samples which can be sectioned.^16^ The removal of the paraffin involves treatment with the aggressive solvent xylene which is very likely to damage the lipidic layer of mycobacteria.^16–18^ Xylene has been implicated in the low sensitivity of AFS, although it was erroneously believed that MA were the target of AFS. ^10^ This study tries to clarify the target of fluorescent AFS and addresses the detrimental effect of xylene by investigating novel, simple, solvent-free approaches as alternatives to xylene deparaffinization and their effect on AFS staining.

## Methods

### Smears

*Mycobacterium bovis*, BCG, was cultured in Middlebrook 7H9 medium (Difco, Detroit, MI, USA) at 37°C in 5% CO_2_ atmosphere. Smears from culture (diluted 1:4) were air-dried and fixed with methanol. These smears were used either as untreated controls or subsequently submerged in 100% xylene for 10 minutes (post-fixation treatment). Alternatively, before the preparation of smears, mycobacterial cultures were either pre-incubated with 100% xylene (4:1) for 10 minutes (pre-incubation treatment); or fixed in 10% formalin (1:1), then centrifuged for 5 minutes at 12300 RCF, and the sediment incubated in 10%, 50%, 100% xylene for 20 minutes.

### Paraffin embedded tissues

Formalin-fixed-paraffin-embedded (FFPE) tissue specimens with a diagnosis of infection by mycobacteria were provided by the *Faculdade de Medicina Veterinária da Universidade de Lisboa*. Tissues were sectioned with 4μm thickness. Standard deparaffinization was performed using xylene (100%) for 10 minutes; repeated with clean xylene (100%) for 10 minutes; followed by ethanol 100%, 95%, 70%, each for 5 minutes and a final 5-minute hydration step in water. Sections were also heat-treated in the Lab Vision^™^ PT-Module^™^ (Thermo Fischer Scientific, Waltham, MA, USA), with pH 6 buffer and exposed to 65°C and then 95°C for 20 minutes each. For the Projected-Hot-Air-Deparaffinization (PHAD) method a common hair dryer (SilverCrest® SHTK 2000W B1, Lidl, Neckarsulm, Germany) was set up to project hot air at maximum blowing force, perpendicular to the slide, from 22cm distance, maintaining 72.5°C at the section surface for 20 minutes. Sections were then submerged in distilled water at room temperature for 5 minutes.

The three solvent free methods, (i) water bath, (ii) dry heat oven, and (iii) centrifugation at high humidity and temperature provided poor results and are described in the supplementary information, as are detailed descriptions of the set-up and optimization of the PHAD method (figure S3-S5).

### Staining

Standard Kinyoun and Auramine O staining protocols were used as described elsewhere ^19^ and details can be found in the supplementary information. In brief, fixed smears or tissue section were stained for 15 minutes with Auramine O (0.5g AuO, Merck, Darmstadt, Germany in 50ml of 95% ethanol mixed with 420ml of distilled water containing 15g phenol, AppliChem, Darmstadt, Germany), or 2 minutes with carbolic fuchsin (Ziehl-Neelsen)(4g of basic fuchsin, Sigma-Aldrich, St. Louis, MO, USA, in 20ml of 95% ethanol mixed with 100ml of distilled water containing 8g of phenol). After rinsing, they were decolorized with 0.5% acid-alcohol (0.5ml of hydrochloric acid in 100ml of 70% ethanol). After another rinsing step, they were counterstained for 2 minutes with 1% methylene blue in distilled water (Alfa Aesar, Karlsruhe, Germany). A standard Hematoxylin and Eosin (H&E) staining protocol was used as described elsewhere ^20^ with minor modifications (see supplementary information).

For triple-staining, smears were first stained with Hoechst-33342 working solution (Stock solution was prepared by dissolving 5.62mg of Hoechst-33342 in 10ml of distilled water and diluted 1:20 in distilled water for working solution) for 30 minutes. After rinsing in tap water, they were stained with 50μM Pyronin Y working solution (Stock solution was prepared by dissolving 3.03mg of Pyronin Y in 10ml of distilled water and diluted 1:20 in distilled water for working solution). After rinsing, they were stained with the described AuO staining protocol.

### Microscopy

Optical and fluorescence photographs were obtained using a DM2500 epifluorescence microscope (Leica Microsystems – Wentzler, Germany) or a confocal laser point-scanning microscope (Zeiss LSM 880, Germany) (details in supplementary information) with the respective optical set-up for imaging the fluorescence of each respective dye. For the AuO stain this consisted in the Leica filter cube L5: excitation BP-filter 480/40nm, dichroic mirror 505nm, suppression BP-filter 527/30nm.

For fluorescence measurements, 20 photographs were taken of different fields with a Leica DFC480 camera using FireCam software. Camera settings were standardized for sample/stain type to allow comparisons (see supporting information). Image J (version 1.52a) software was used to measure fluorescence of individual bacilli (clumps, cords or overlapping cells were ignored).

### Statistical analysis

All analyses were performed using R Statistical Software version 4.0.4 (2021-02-15). Shapiro-Wilk normality test confirmed the non-normal nature of the fluorescence measurements data and thus the non-parametric Wilcoxon ranked-sum test was used. These tests and corresponding confidence intervals for the difference in median value was calculated with the R package *stats* version 4.0.4. The effect size estimate is expressed as the *r*-value of correlation coefficient^21^, including 95% confidence intervals as margin of error, calculated with the *rstatix* version 0.7.0.

## Results

Using the standard AuO staining method, mycobacteria were clearly visible in smears and in tissue sections (figure 1). Higher magnification of the smears showed an intracellular beaded staining pattern of mycobacteria, which was also visible in tissue sections (figure 1A and 1B), allowing to distinguish them easily from other potentially fluorescent artefacts. Importantly, little unspecific background fluorescence of tissue was noted. Specific DNA (Hoechst-33342) and RNA (Pyronin Y) stains colocalized with the acid-fast stain AuO inside the mycobacteria (figure 1C). The intracellular staining was heterogeneous, with beaded pattern.

**Figure 1.**
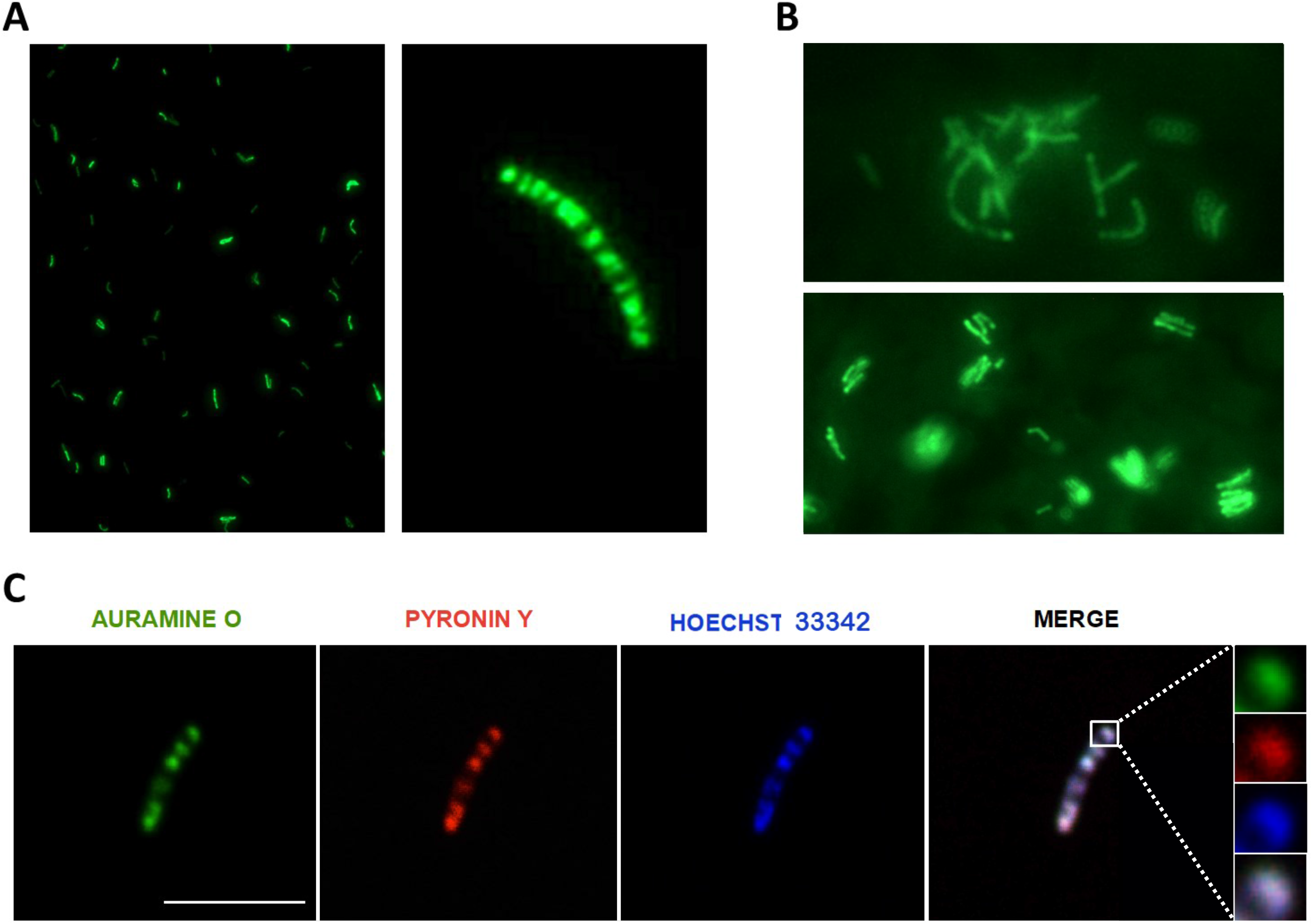
Colocalization of Auramine O with DNA and RNA specific stains in *M. bovis*. *M. bovis* in liquid culture (A) and mycobacteria in tissue from feline dermis (B) show bright fluorescence with a typical inhomogeneous, beaded pattern at higher magnification and little to no cell wall staining. Triple staining of cultured *M. bovis* with AuO and the RNA-specific stain Pyronin Y and the DNA-specific stain Hoechst-33342 shows almost perfect colocalization (C) which suggests that the target of AuO are nucleic acids. In prokaryotes these are organized in nucleoids, which is compatible with the inhomogeneous, beaded pattern (A, B). Fluorescent microscopy, Amplifications: (A) 200x & 1000x, (B)1000x (C) 1000x, scale bar: 5μm.

*M. bovis* (BCG) grown in liquid culture showed reduced fluorescence compared to untreated control (median = 50.5, n = 1128 observations) when exposed to 20% xylene, both post-fixation (median = 33.0, p<0.0001, n = 625 observations, 95% CI of difference: 14.99 - 19.00, *r* = 0.33 ± 0.04, moderate effect size) or when pre-incubated (median = 36.0, p<0.0001, n = 2415 observations, 95% CI of difference: 11.99 - 15.00, *r* = 0.28 ± 0.03, small effect size). *M. bovis* BCG grown in liquid media displayed a typical cording which was easily visible after staining with the standard AuO method (figure S2).^22–24^ Interestingly, preincubation with different xylene concentrations (10%, 50% and 100%) produced images where cord-like structures were seen to increasingly disintegrate into pools of individual mycobacteria (figure S3).

The results of several heat-based approaches which were tested to remove the paraffin, which melts at above 50°C, showed that three methods were ineffective in removing paraffin for subsequent staining: (i) the formalin-fixed paraffin-embedded tissue sections overnight at 68°C in a dry heat oven, (ii) in a water bath at 72°C, and (iii) centrifuging the slides in a high humidity / high temperature atmosphere. Contrary to this, the projected hot air deparaffinization (PHAD) allowed successful staining.

Microscopic examination of tissue sections deparaffinized with PHAD showed a significant increase in the number and fluorescence intensity of mycobacteria when compared to the tissue sections deparaffinized with the standard method, using xylene (figure 2). Given the apparent superiority of PHAD, the method was further characterized, defining optimal running time, temperature profile and standardized distances of the heat source (hair dryer) to the sample (figures S3-5, supplementary file).

**Figure 2.**
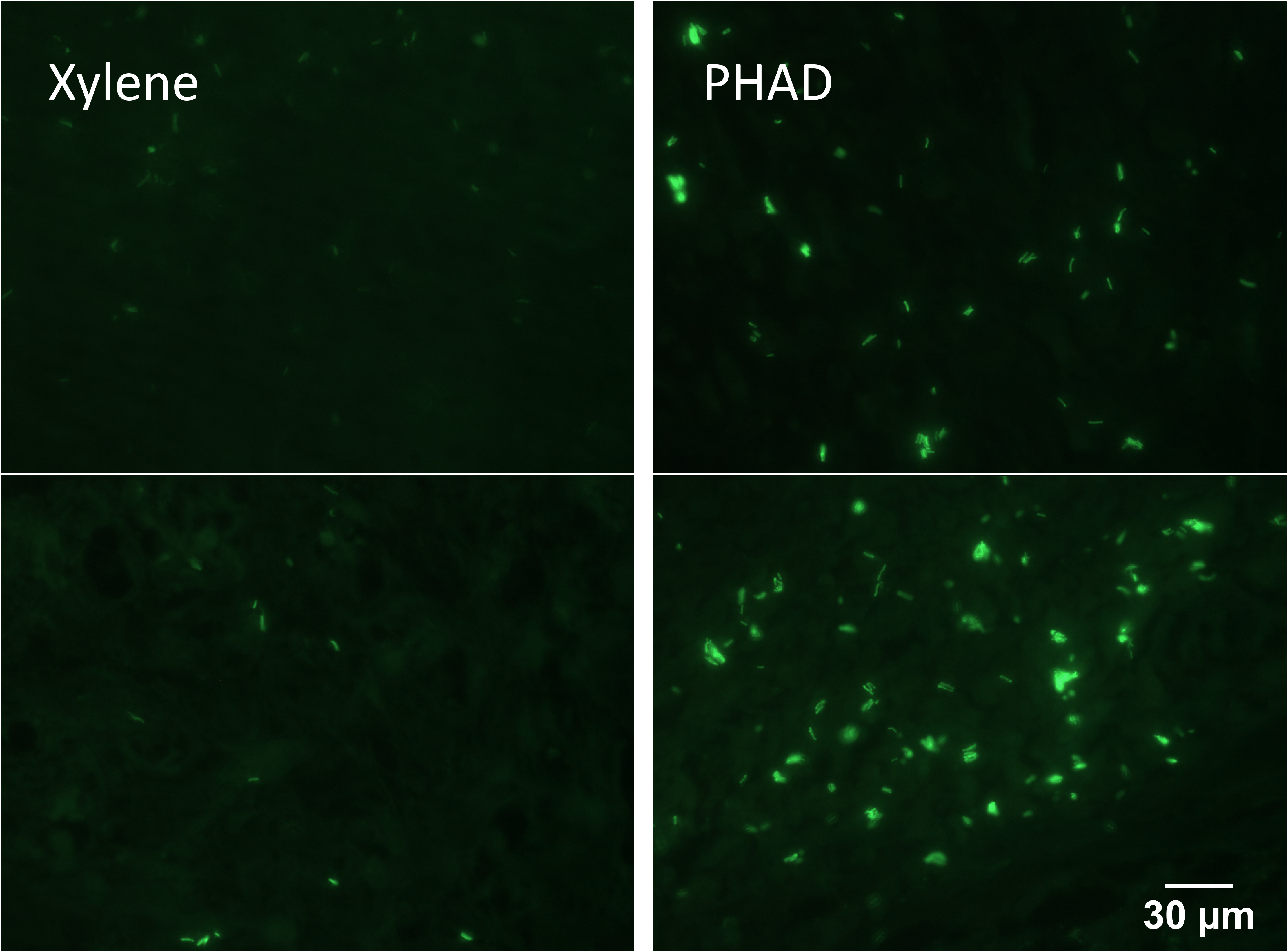
Representative image of AuO stained section after xylene and PHAD deparaffinization. Mycobacteria infected muscle tissue stained with the AuO method. The images show the fluorescence after standard xylene deparaffinization (left) and after Projected Hot Air Deparaffinization (PHAD) (right). (1000x).

Tissue integrity after PHAD deparaffinization was confirmed by observation of H&E-stained sections (figure 3, top panel). Cell structures and tissue organization appeared unchanged comparing the PHAD and standard xylene method. However, H&E staining after PHAD revealed areas of H&E staining with some variation in stain intensity, indicating that the uneven deparaffinization prevented the dyes from fully penetrating the tissues. This did not occur with the Ziehl-Neelsen (ZN) staining, which was intense and uniform. Most importantly, the number of mycobacteria in PHAD treated sections was significantly higher (figure 3, bottom panel).

**Figure 3.**
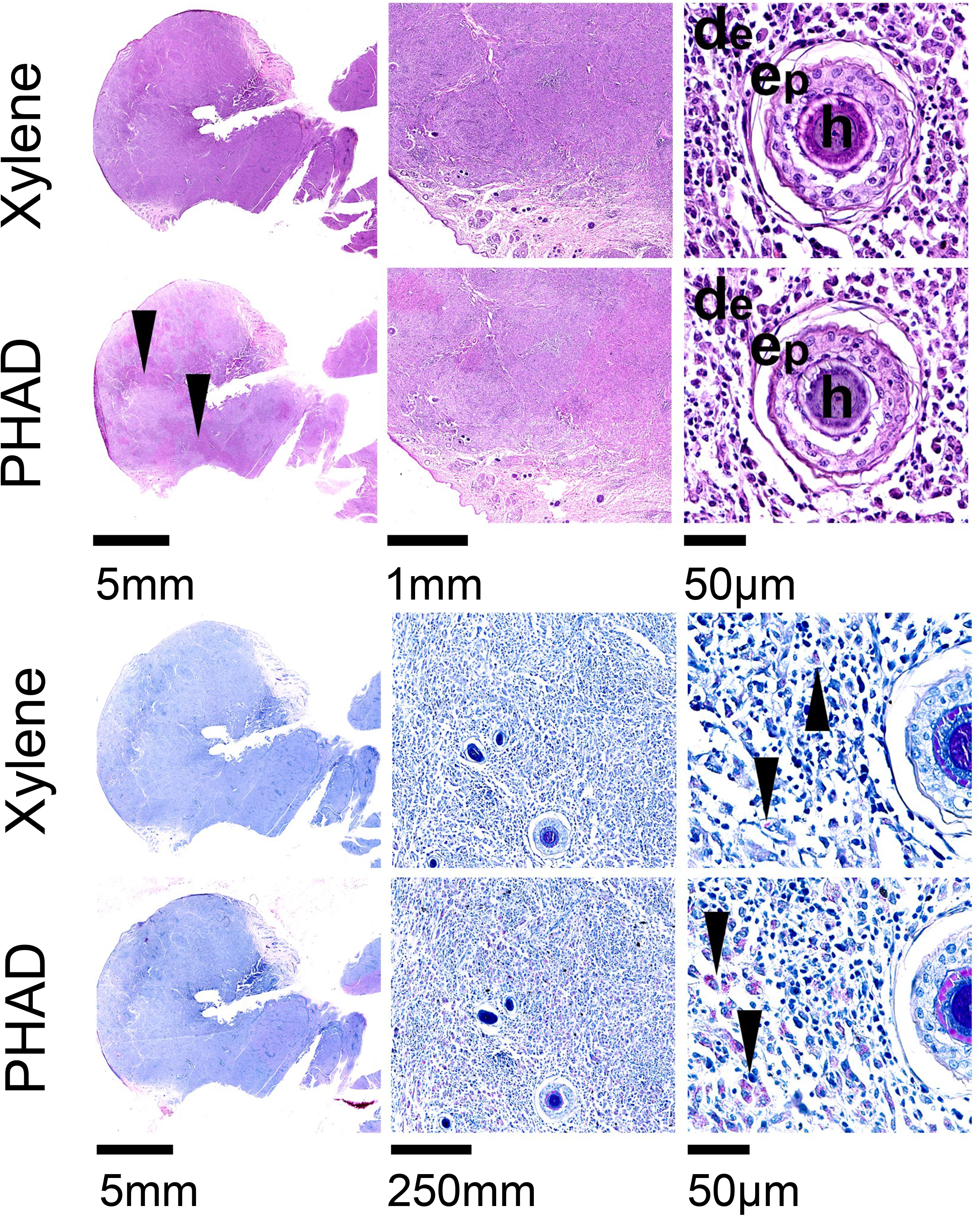
Tissue integrity and mycobacterial yield after PHAD and xylene treatment. Tissue integrity and morphology is maintained as shown in H&E-stained sections of skin (top panel), although some light-coloured patches indicate unevenly stained sections (arrowheads), likely due to incomplete deparaffinization with the PHAD method (Magnifications: 5x, 2.5x and 40x). Ziehl-Neelsen-stained sections show significantly more acid-fast bacilli (arrowhead) with the PHAD method as compared to the xylene deparaffinization (bottom panel) (Magnifications: 5x, 10x and 40x, de: dermis, ep: epidermis, h: hair).

Finally, the performance of the PHAD method was compared with xylene deparaffinization, and the Lab Vision^™^ PT-Module^™^, a method used in immunohistochemistry which relies on a heated bath with proprietary buffers. The PHAD showed significantly more mycobacteria and an increased fluorescence in intestinal lymph node and muscle tissue samples (figure 4). While the PT-Module^™^ was superior to the standard xylene method, it performed significantly worse than the PHAD method.

**Figure 4.**
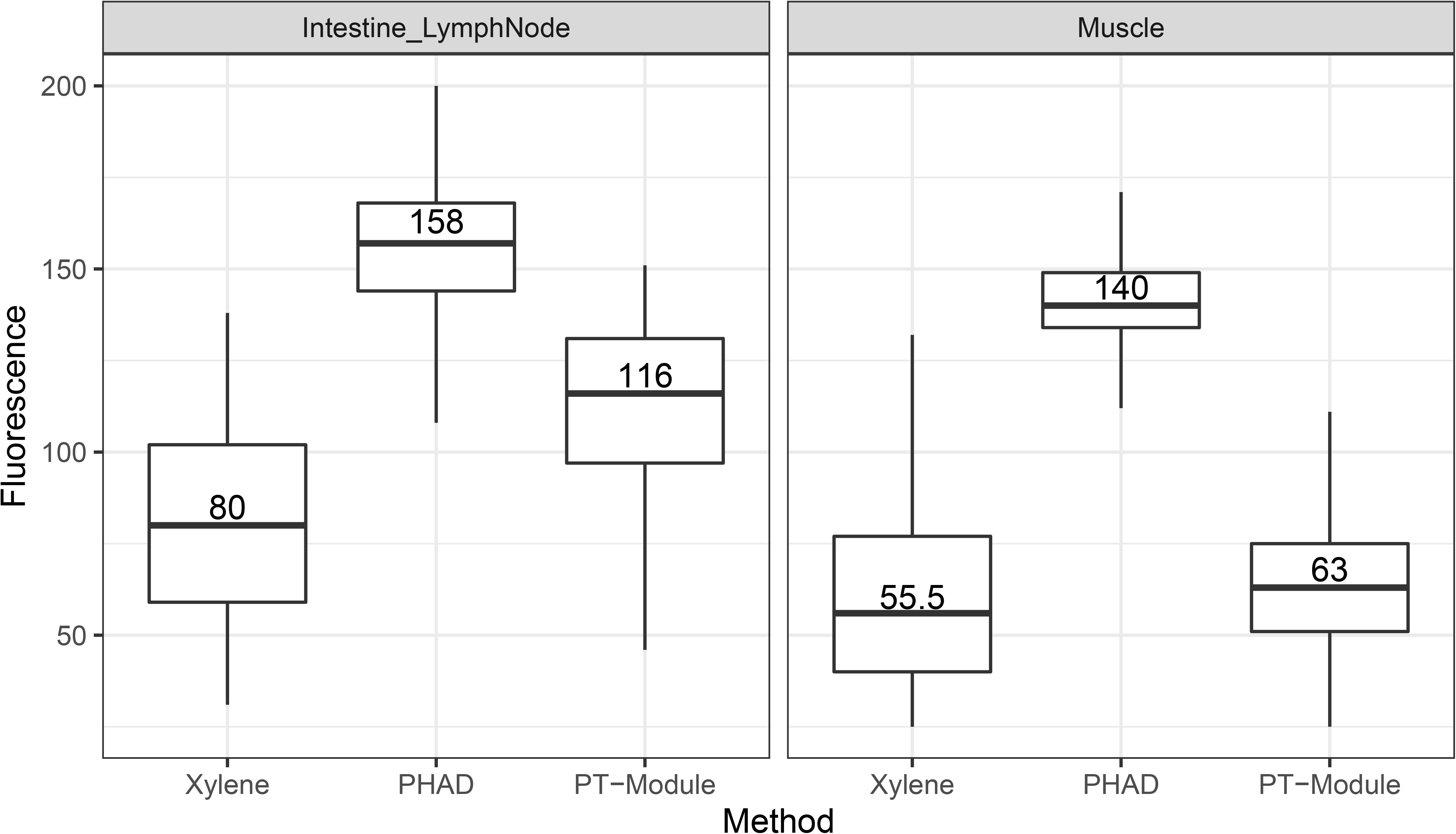
Mycobacteria in tissue fluoresce significantly more after PHAD treatment. Mycobacteria in lymph-node and muscle fluoresce significantly less after standard xylene deparaffinization. While the proprietary PT-Module^™^ increases fluorescence, the solvent free deparaffinization (PHAD) produced the strongest fluorescence. The box represents the values from the first to third quartile, with a horizontal line marking the second quartile/median. The whiskers extend to the minimum and maximum values. Number of bacteria observed in lymph node (LN): n_xylene_ = 86, n_PHAD_ = 882, n_PT-Module_=1201; and muscle (M): n_xylene_=1096, n_PHAD_=1187, n_PT-Module_=1418. LN Xylene - PHAD: p<0.0001.95% CI of difference: 71.9 - 84.0, *r:* 0.48 ± 0.05 (moderate effect size). LN Xylene - PT-Module^™^: p<0.0001, 95% CI of difference 26.0 – 38.0, *r:* 0.24 ± 0.05 (small effect size). M Xylene - PHAD: p<0.0001, 95% CI of difference: 81.9 - 85.0, *r:* 0.85 ± 0.005 (large effect size). M Xylene - PT-Module^™^: p<0.0001, 95% CI of difference: 4.9 - 8.00, *r:* 0.14 ± 0.04 (small effect size).

## Discussion

Extrapulmonary tuberculosis (EPTB) is notoriously difficult to diagnose with microbiological methods.(3) Biological samples are often biopsy specimens containing few mycobacteria.^3^ In those cases, histology aids significantly in establishing a diagnosis, although often only unspecific changes are noted, such as granulomas. For a definitive diagnosis the demonstration of acid-fast bacteria is necessary, usually done by ZN staining, which is well known to be often negative.^3,5,6^

Interestingly fluorescent AFS, such as AuO, which are superior to ZN and have been endorsed by the World Health Organization (WHO) for microbiology diagnosis are not widely used in histological diagnosis.^25^ The results of this study show that mycobacteria in tissues are highly fluorescent after staining with standard AuO with acceptable background fluorescence in tissues (figure 1). Many sources propose that the target of AFS are mycolic acids,^10–12^ which, if true, should result in homogenously stained bacilli. However, what is observed is the extremely typical morphology of mycobacteria (beaded pattern) which makes it easy to distinguish them from artefacts. Fuchsin has been an important chromosome dye since the 1930s, ^26,27^ and fluorescent microscopy of ZN-stained smears (excitation: green – emission: red) reveals the same beaded appearance.^28^ Auramine O was used in the first commercial reticulocyte counter to stain nucleic acids.^29^ Our colocalization results provide strong evidence that the main target of AFS are indeed nucleic acids inside the mycobacteria (figure 1). Interestingly, the typical beading could thus be explained with the distribution of nucleic acids in procaryotes, which cluster in nucleoids. ^30^

The implication of these results is that the performance of acid-fast-stains for diagnosis is dependent on the integrity of the MA rich cell-wall to avoid dye removal during the acid-alcohol differentiation step. In histology two processing steps could have largely detrimental effects for the staining and identification of bacteria: 1^st^ the sectioning of tissue, and 2^nd^ the deparaffinization technique.

Contrary to microbiological smears, where the intact biological sample is smeared on a glass-slide, in histology the tissue is cut in thin (3-4μm thick) sections.^16^ Assuming that mycobacteria are around 0.2–0.5μm wide and 2–4μm long and are heterogeneously distributed and oriented in the tissue, many of them will also be sectioned during this process. This exposes the intracellular contents to the decolorization step, causing bacilli to lose their acid-fastness. In fact, Koch himself observed that grinding mycobacteria reduced their acid-fastness.^31^

Perhaps much more important is the processing of the tissue sample. Contrary to what was previously reported in one study^10^, we saw no difference between formalin fixed or unfixed culture smears, nor any cord disintegration in formalin fixed cultures (results not shown). Furthermore, many research applications use nucleic acid stains successfully in formaldehyde fixed and deactivated cells and tissues.^32^

Xylene on the other hand is a potent organic solvent and one can expect it to damage the MA layer of mycobacteria and render them less fluorescent. Our results strongly confirm this idea and extend previous reports^10^ by using AuO and a method to measure and quantify fluorescence objectively. By observing a dose-dependent effect on the disintegration of cords, xylene seems to damage the MA rich extracellular matrix which is responsible for the phenomenon of cord-formation. ^22,23^

The strong reduction of fluorescence of xylene-exposed mycobacteria, in smears but also in tissue, is further evidence of the deleterious effect of xylene on acid-fast-staining.

To address this, a completely solvent-free deparaffinization method was established, based on dry heat. Using a simple hair dryer, the PHAD proved to be highly effective in removing the paraffin to allow acid-fast staining. The mechanism likely depends on two factors: first, the effective and consistent application of dry heat to melt paraffin; and second, the strong air current projected onto the slide removes the liquid paraffin from the tissue. H&E staining confirmed that tissue integrity as well as morphology are maintained. Nonetheless, the PHAD deparaffinization seems to be somewhat incomplete and uneven as indicated by some variation in the H&E staining and may not be suitable for all staining techniques.

ZN-staining seems unaffected and allowed easy identification of acid-fast bacteria. Of crucial importance, the number of mycobacteria present in the tissue appears to be much higher after PHAD processing, which suggests that this method could increase the sensitivity in paucibacillary cases. Quantitative assessment of this effect after AuO staining corroborated these observations, with significantly stronger fluorescence of mycobacteria (large effect sizes) in PHAD, comparing to the standard deparaffinization with xylene. Of note, PHAD outperforms another commercial system using heat, but which also employs a citrate buffer. Interestingly, the results are also in line with older reports which described a reduction in detectible auramine-rhodamine-stained bacilli after pure xylene treatment of tissue, which was mitigated once the solution was replaced with a mixture of xylene and ground-nut oil.^33^ The higher sensitivity of the Fite stain, in comparison to ZN, provides further support, since it uses less xylene in the deparaffinization and mounting process.^10,34^

In conclusion, AuO is a fluorescent nucleic acid dye which can be used to identify acid-fast mycobacteria in tissues. Both AuO and ZN stains highlight the typical morphology of beaded mycobacteria. Xylene, used in standard deparaffinization methods, also seems to damage the mycobacterial cell-wall, crucial for AFS, and consequently reduces acid-fastness. The performance of AFS in histology could be significantly enhanced if sections are deparaffinized with solvent-free methods, namely dry heat, as the results of this research suggest. This might have the potential to improve the histological diagnosis of tuberculosis significantly.

## Funding

No funding was received for this project and all experiments were carried out with already existing materials and reagents.

## Disclosure

The authors declare no conflicts of interest.

## Contributions

Pedro Marinho designed and performed the experiments, analyzed the data, and wrote the draft manuscript. Soraia Vieira performed the triple staining experiments and contributed to the draft manuscript. Tânia Carvalho and Maria C. Peleteiro gave expertise input on the histology aspects, provided samples, and evaluated the H&E and ZN-stained sections. Thomas Hanscheid designed the overall study, advised on experimental design, assisted in data analysis, and contributed to writing the final manuscript.

## Notes

### Competing Interest Statement

The authors have declared no competing interest.

