## Supplementary Information and Figures for "A novel and simple heat-based method eliminates the highly detrimental effect of xylene deparaffinization on acid-fast stains"

**Running head:** Negative effect of xylene on acid-fast-stains

**Methods:**

Reagents: Auramine O solution (3.3mM) was prepared by dissolving 0.5g of Auramine O powder (Merk – Darmstadt, Germany) in 50ml of 95%(v/v) ethanol. This is then mixed in a solution with 15g of phenol crystalline (AppliChem – Darmstadt, Germany) dissolved in 420ml of distilled water. The flask is kept in the dark at room temperature (RT). Carbol-Fuchsin (9.2x10-2μM) was prepared by dissolving 4g of Basic Fuchsin powder (Sigma-Aldrich – St. Louis, MO, USA) in 20ml of 95%(v/v) ethanol. This is then mixed in a solution with 8g of phenol crystalline dissolved in 100ml of distilled water. The flask is kept in the dark at RT. Decolorizing acid-alcohol solution 0.5%(v/v) was prepared by adding 0.5ml of concentrated hydrochloric acid (HCl) to 100ml of 70% ethanol(v/v). Counterstain Methylene Blue 1%(w/v) (Alfa Aesar – Karlsruhe, Germany) was prepared by dissolving 2g in 200ml of distilled water each. Hoechst-33342 solution (1mM) was prepared by dissolving 5.62mg of Hoechst-33342 powder in 10ml of distilled water (Sigma-Aldrich – St. Louis, MO, USA). The working solution with a concentration of 50μM was obtained by diluting the stock solution 1:20 in distilled water. Pyronin Y stock solution (1mM) was prepared by dissolving 3.03mg of Pyronin Y powder in 10ml of distilled water (Polysciences – Warrington, PA, USA). The working solution (50μM) was prepared by diluting the stock solution 1:20 in distilled water. The Hoechst-33342 and Pyronin Y stock and working solutions were stored at 4ºC in the dark.

Auramine O and Kinyoun (ZN) staining: The smear/section is flooded with the Auramine O or carbolic-fuchsin solution for 15 minutes or 2 minutes respectively, and then rinsed gently with running tap water applied on the edge of the slide and never on top of the sample. Then the decolorizing solution is applied drop by drop until no more color is visible on the smear. Again, the smear is rinsed and then flooded for 2 minutes with the counterstain, rinsed with tap water, and left to air-dry.

Hematoxylin and Eosin Staining Protocol (H&E): slides are flooded with Harris Hematoxylin (Bio-Optica – Milano, Italy) for 3 minutes, washed with running water for 5 minutes, applied not directly to the sample; they are then dipped once in 70% ethanol and four times in alcoholic eosin (Sigma-Aldrich – St. Louis, MO, USA). Finally, slides are flooded with 70% ethanol for 30 seconds, 95% ethanol for 30 seconds, 100% ethanol for 30 seconds and again with clean 100% ethanol. Slides are immediately mounted as explained below.

Hoechst 33342, Pyronin Y and Auramine O triple-staining: Smears were first stained with Hoechst-33342 working solution for about 30 minutes. The slides were then gently rinsed with tap water and flooded with Pyronin Y working solution for 30 minutes. The slides were then rinsed with water and stained with the Auramine O staining protocol as described. Finally, the triple stained smears were air-dried and mounted. The slides were covered with an aluminum foil during and after the staining procedure.

Mounting and observation: Glass slides with tissue sections were mounted with a coverslip for observation using BacLight^TM^ mounting oil (Invitrogen – Eugene, OR, USA). For permanent mounts, tissues were dehydrated as described in the final steps of the H&E method above or left to air dry completely if using the ZN stain; they were flooded with xylene (Leica Biosystems – Nussloch, Germany) for 10 minutes and then two drops of Quick-D mounting medium (Klinipath – Duiven, Netherlands) were put on the edges of the glass slide, the coverslips were dipped in xylene and placed carefully on top of it. Air bubbles were gently pushed out and the mounted slide was dipped in xylene to clean excess mounting medium. The slides were left for 24h at RT to dry and solidify. Except for the triple-stain slides, smear samples were observed directly, not mounted.

Imaging: Optical and fluorescence photographs were obtained using a DM2500 epifluorescence microscope (Leica Microsystems – Wetzlar, Germany) with the following specifications: 100W mercury lamp,

For Hoechst-3342 fluorescence (ultraviolet-blue) the A4 filter cube was used: excitation BP-filter 360/40nm, dichroic mirror 400nm, suppression BP-filter 470/40nm, for AuO fluorescence (blue-green) the filter cube L5 was used: excitation BP-filter 480/40nm, dichroic mirror 505nm, suppression BP-filter 527/30nm), and for Pyronin Y (green-red) the filter cube N2.1 was used: excitation BP-filter 515-560nm, dichroic mirror 580nm, suppression LP-filter 590nm) was used.

The DFC480 camera (Leica Microsystems – Wetzlar, Germany) parameters used for photographs taken for posterior fluorescence measurements, both for tissue sections and smears stained with AuO, were standardized in the Leica FireCam software. Smears: Exposure - 580ms, Gain - 1x, Levels - 0; 2.0; 223, Color - Light Red, Saturation - 80%. Tissue: Exposure – 1.7s, Gain – 1.7x, Levels – 0; 2.0; 223, Color – Light Red, Saturation – 80%.

Confocal images were acquired with a confocal laser point-scanning microscope (Zeiss LSM 880, Germany) and the ZEN 2.1 software (black edition, Zeiss, Germany) using gain settings between 600 and 750, a scan speed of seven and a mean of two scans per pixel. The frame-size was used according to the optimal values suggested by the software. Zoom function was set according to the organism size and the pinhole was set to one airy unit. To achieve higher resolution and an improved signal-to-noise, the Airyscan "27 system was used to acquire all the confocal images. The lasers used for confocal microscopy are the following: Hoechst-33342 - Diode 405-30 (405nm); Auramine O - Argon (488nm); Pyronin Y - HeNe 594 (594nm).

PHAD standardization: The hairdryer was fixed in a grid, where the nozzle of the hairdryer was at 21cm from the benchtop and 22cm from the FFPE slide in a perpendicular position. All components were taped down and static for all experiments. Temperature was measured with a mercury thermometer every 2 minutes, for 20 minutes plus additional measurements after shutting off the hair dryer and submerging in distilled water. The thermometer was taped to the same support as used for FFPE slides and the hairdryer was set to the maximum temperature. Air speed could not be determined, but maximum setting was also used. Duration of the process was chosen by conducting the deparaffinization at several different periods of time: 10, 20, 30 and 40 minutes (shorter times were not tested because they proved inefficient in initial attempts). It was found that between 10 and 20 minutes no further difference could be seen, although 20 minutes showed a slightly higher median value and appeared to reveal more bacilli.

Additional deparaffinization methods: For the dry heat oven method sections were placed vertically at 68ºC overnight, then placed in distilled water for 5 minutes immediately followed by staining. Water bath deparaffinization consisted in the submersion of the tissue sections in distilled water, each in individual containers, for a duration of 20 minutes, whose temperature was kept at around 72ºC. After this time, the slides were immersed in distilled water at RT for 5 minutes immediately followed by staining. Centrifugation at high humidity and high temperature consisted in fixing glass slides vertically, facing outwards, on a rotor of a centrifuge of a domestic appliance; the rotor is placed inside an enclosed chamber. Water was heated to boiling point and poured into the chamber to about 1cm below the rotor. The centrifuge is immediately turned on for a cycle lasting 5 minutes which reached 68ºC of air temperature. At the end of the cycle, the glass slides are submerged in distilled water at RT for about 5 minutes.

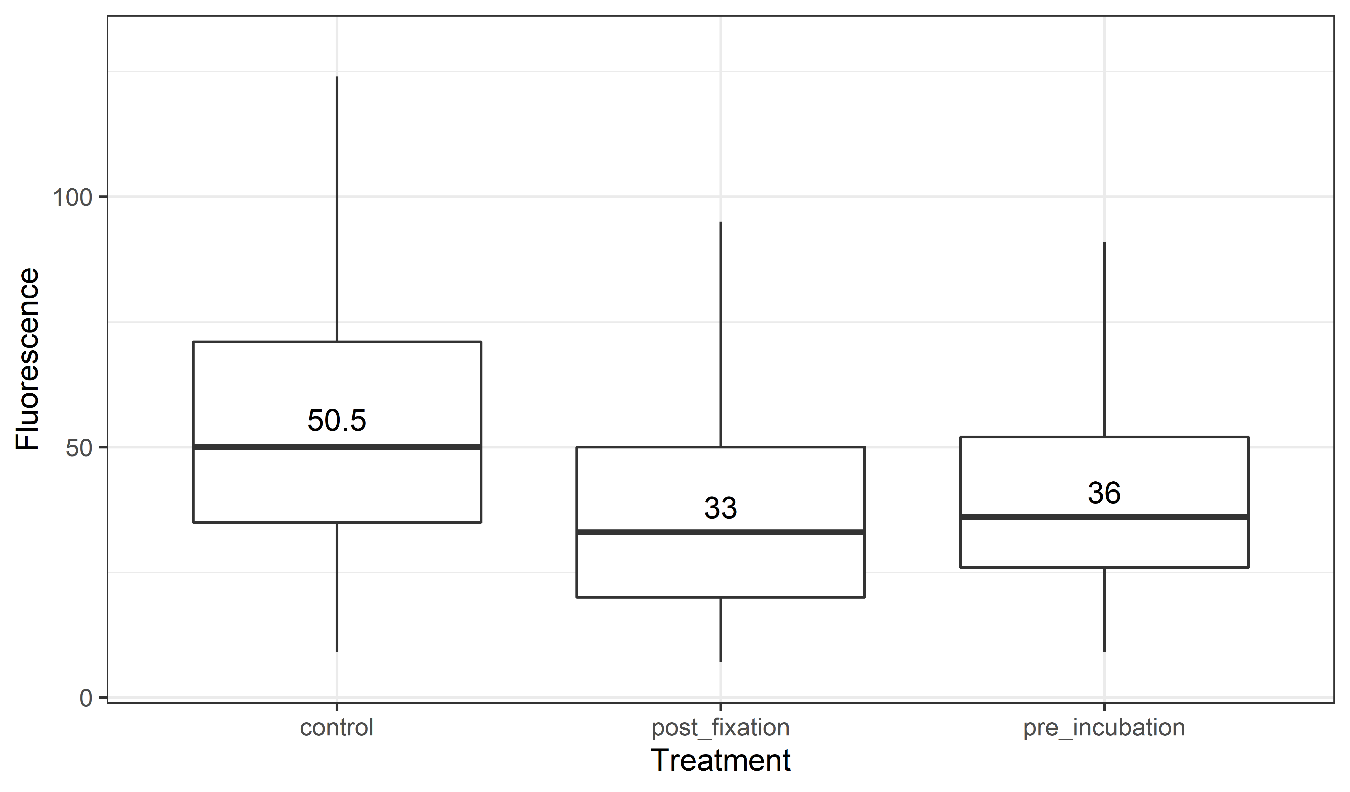

**Figure S1 – M. bovis (BCG) fluorescence is reduced after xylene exposure**

*M. bovis* (BCG) grown in liquid culture, exposed to 20% (v/v) xylene showed reduced fluorescence, both post-fixation and when preincubated as compared to untreated control. The box represents the values from the first to third quartile, with a horizontal line marking the second quartile/median. The whiskers extend to the minimum and maximum values. Number of observations: n_post_ =625, n_pre_ = 2415, n_control_ = 1128; post-fixation: p<0.0001, 95% CI of difference: 14.99 - 19.00, *r*: 0.33 ± 0.04 (moderate effect size); preincubation: p<0.0001, 95% CI of difference: 11.99 - 15.00, *r*: 0.28 ± 0.03 (small effect size).

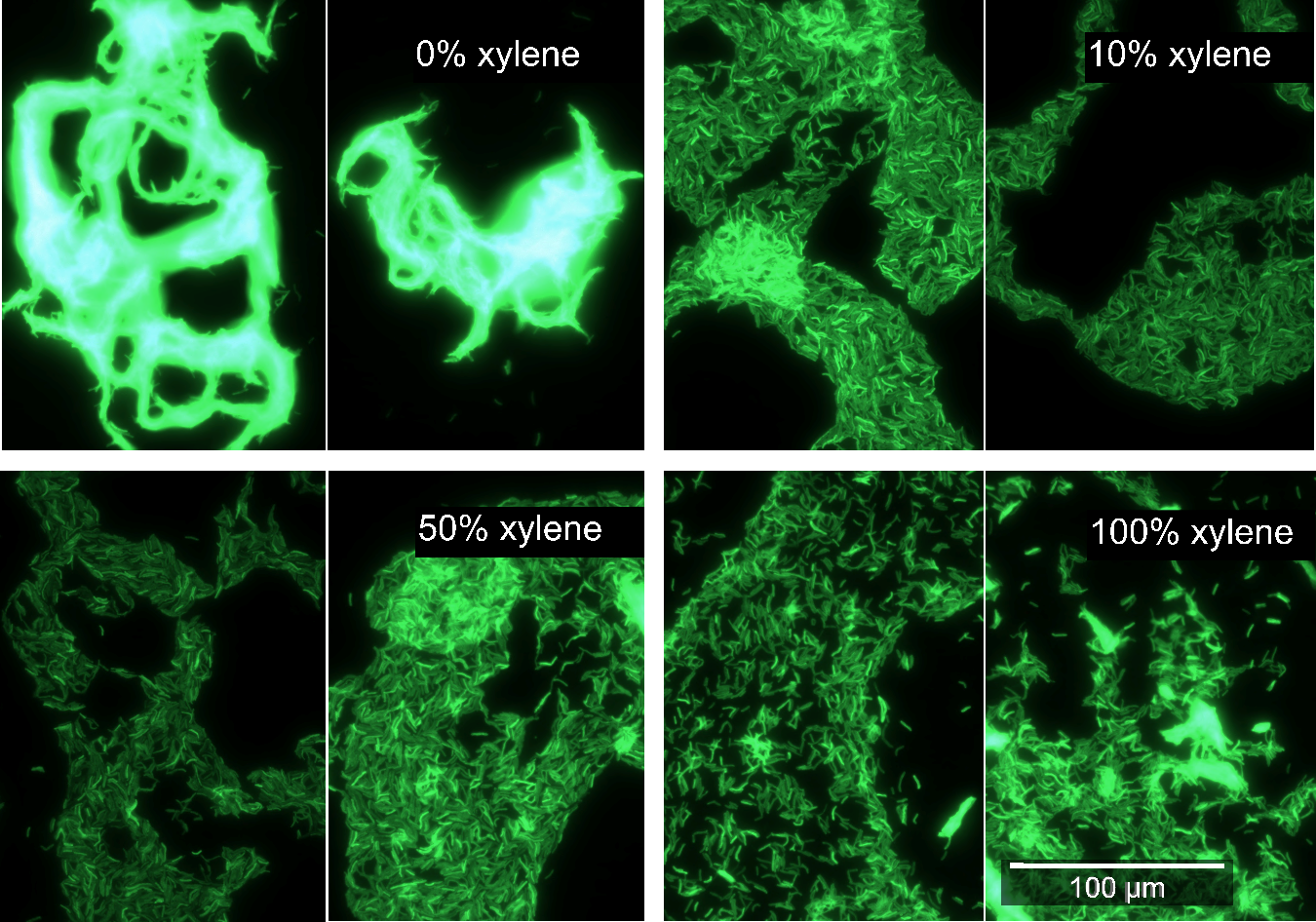
**Figure S2 – AuO staining of *M. bovis* BCG in liquid culture** **preincubated with xylene**

*M. bovis* (BCG) showing typical cording when grown in liquid culture (top-left panel). Preincubation with increasing concentrations of xylene leads to a dose-dependent disaggregation of the cords (top-right and bottom panels). Of note, many individual bacteria were released form the cords, often brightly fluorescent. Fluorescent microscopy. (1000x).

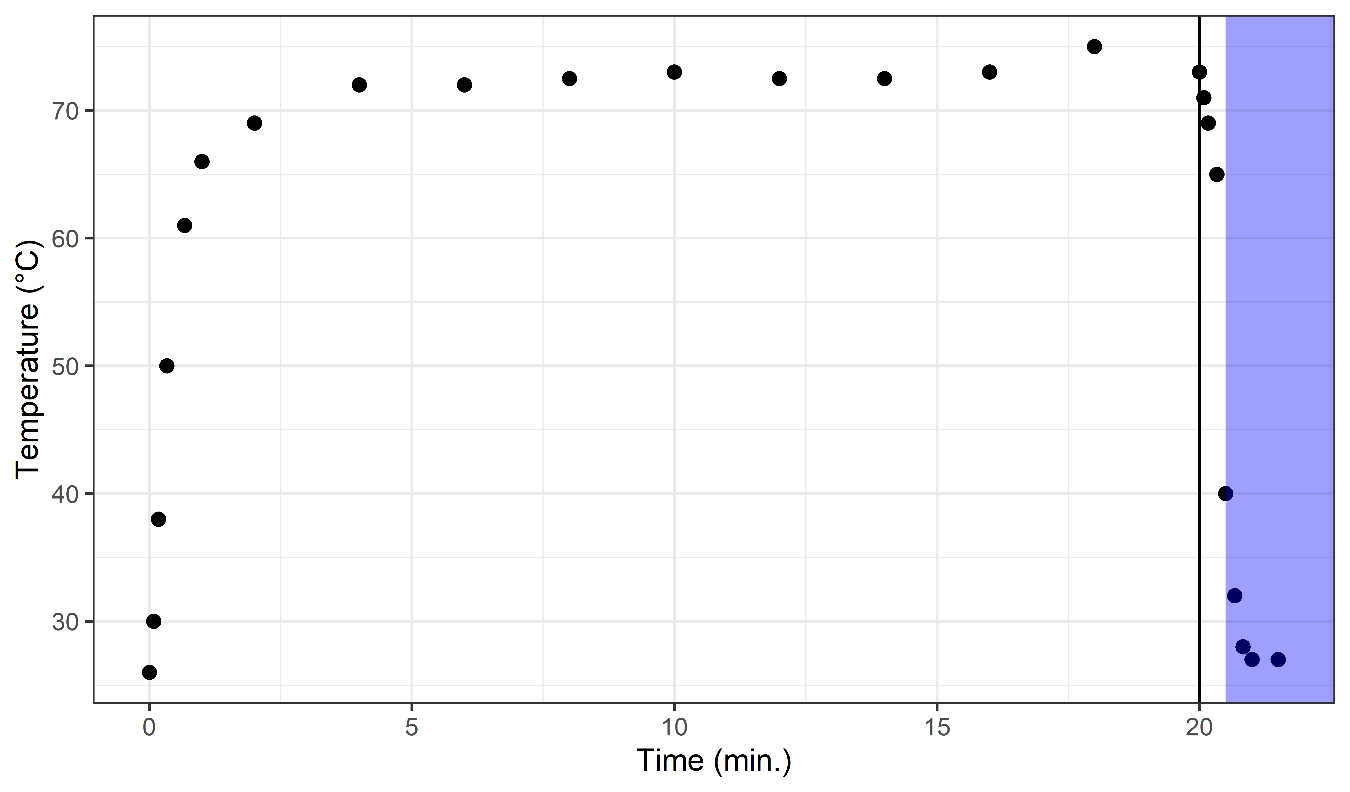
**Figure S3 – Temperature profile of the PHAD method**

Temperature profile observed at the level of the slide under the heat source of the PHAD method. The vertical black line represents the shut-off time point at 20 minutes and the blue rectangle represents the immersion of the sample in water. The temperature during running was approximately 72.5ºC.

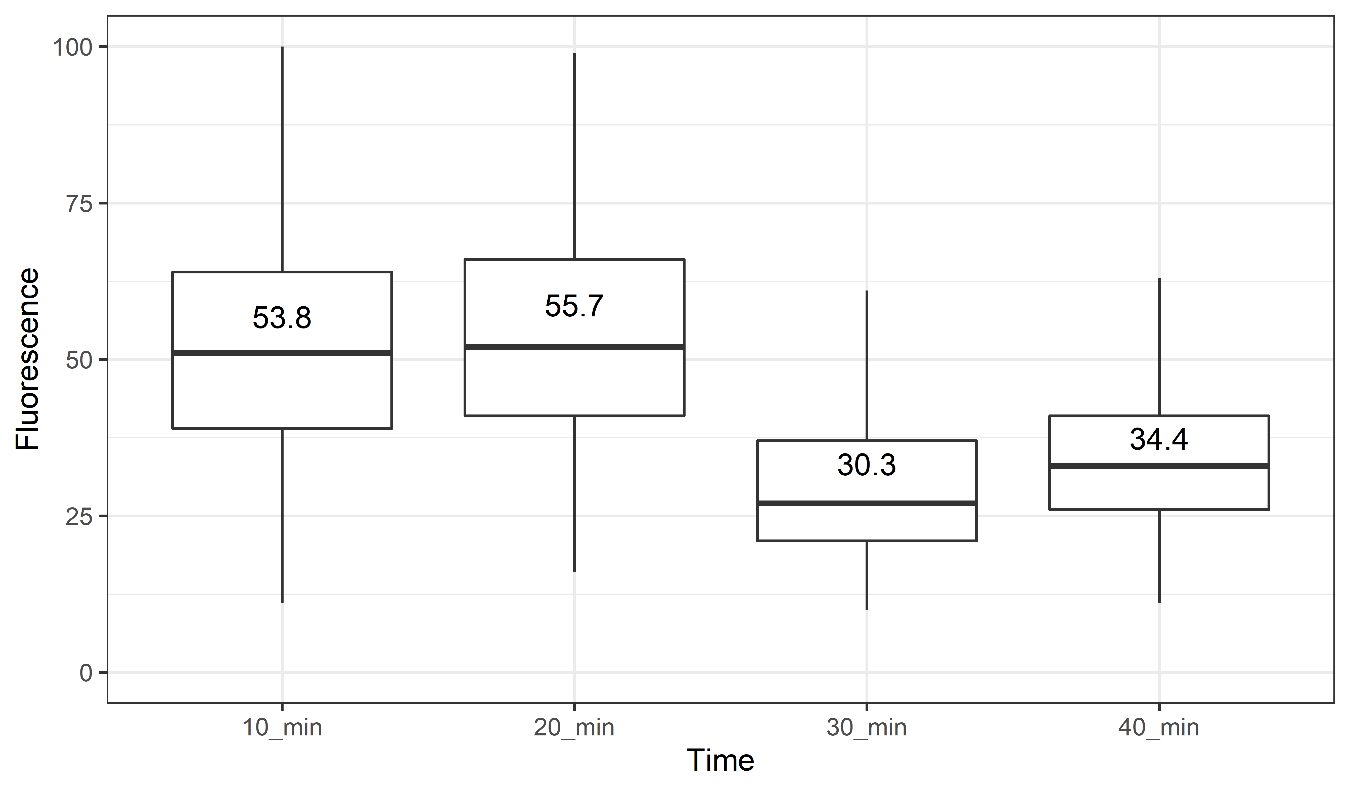
**Figure S4 – Comparison of different application times for the PHAD method**

Comparison of several durations of heat application in the PHAD method, regarding its effect on fluorescence of mycobacteria in bovine muscle tissue sections. The box represents the values from the first to third quartile, with a horizontal line marking the second quartile/median with the corresponding value. The whiskers extend to the minimum and maximum values (number of observations: n_10_=616, n_20_=818, n_30_ =1011, n_40_=461).

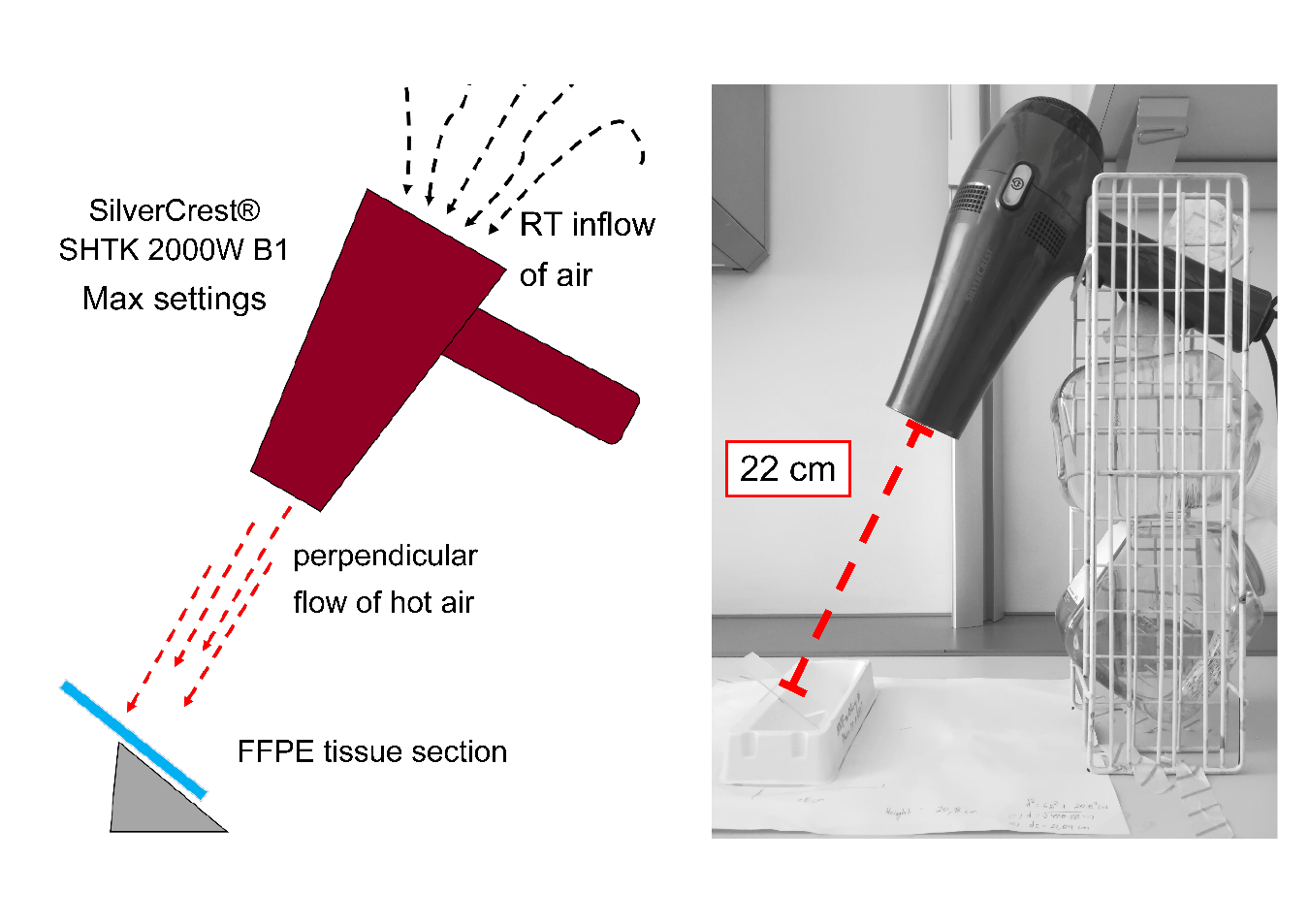
**Figure S5 – Schematic representation and photograph of the PHAD set-up**

A working station was set up with fixed positions for all parts and the sample, to assure reproducibility. On the left is a simple schematic representation of the mechanism and on the right is a photograph of the station and the distance between the source of air and the sample.
